# LIM-HD transcription factors are required for regeneration of neuronal and intestinal cell subtypes in planarians

**DOI:** 10.1101/2023.02.07.527492

**Authors:** M. Dolores Molina, Dema Abduljabbar, Susanna Fraguas, Francesc Cebrià

## Abstract

Adult planarians can regenerate the gut, eyes, and even a functional brain in just a few days after injury. Proper regeneration of these complex structures requires that signals guide and restrict the commitment of their adult stem cells and ensure the identity and patterning of the newly formed structures. During embryogenesis of both vertebrates and invertebrates, LIM Homeodomain (LIM-HD) transcription factors act in a combinatorial ‘LIM code’ that controls crucial aspects of cell fate determination and cell differentiation, including specification of neuronal cell type identity and axonal guidance. So far, however, our understanding about the role these genes may play during regeneration is limited. Here, we report the identification and functional characterization of the full repertoire of LIM-HD genes in *Schmidtea mediterranea*. We found that these *lim homeobox genes* (*lhx*) appear mainly expressed in complementary patterns along the cephalic ganglia and digestive system of the planarian. By functional RNAi based analysis we have identified that several *Smed-lhx* genes (*islet1*, *lhx1/5-1*, *lhx2/9-3*, *lhx6/8*, *lmx1a/b-2* and *lmx1a/b-3*) are essential to pattern and size the planarian brain as well as for correct regeneration of specific subpopulations of dopaminergic, serotonergic, GABAergic and cholinergic neurons, while others (*Smed-lhx1/5.2* and *Smed-lhx2/9.2*) are required for the proper expression of diverse intestinal cell type markers, specifically the goblet subtype. LIM-HD are also involved in the control of axonal pathfinding (*lhx6/8*), axial patterning (*islet1* and *lmx1a/b-3*), head/body proportions (*islet2*) and stem cell proliferation (*lhx3/4*, *lhx2/9-3*, *lmx1a/b-2* and *lmx1a/b-3*) in planarians. Altogether, our results suggest that planarian LIM-HD could provide a combinatorial LIM code to control axial patterning, axonal growing as well as to specify distinct neuronal and intestinal cell identities during regeneration.

## 1. Introduction

Planarians can regenerate damaged or missing tissues and organs or entire organisms to full function within a period of a few days to several weeks. The extraordinary regenerative capacity of planarian relies on the presence of a population of adult stem cells named neoblasts (reviewed in Reddien, 2018; Reddien, 2022; Wagner et al., 2011; Zhu and Pearson, 2016). How neoblasts achieve their final differentiation into the multiple cell lineages remains to be fully answered, although important progresses have been made in recent years (reviewed in Molina and Cebrià, 2021). In any case a combination of intrinsic and external signals and stimuli emanating from differentiated tissues and organs play a role in regulating neoblast biology (Barberán et al., 2016a; Chan et al., 2021; Rossi and Salvetti, 2019; Witchley et al., 2013; Wong et al., 2022) and need to be tightly coordinated and controlled for a successful regeneration.

Transcription factors (TFs) play key roles in multiple aspects of animal development. In planarians, many conserved TFs have been identified (Suzuki-Horiuchi et al., 2021) and characterized to be required for the regeneration of multiple cells, tissues and organs including the intestine (Flores et al., 2016; Forsthoefel et al., 2012; Forsthoefel et al., 2020; González-Sastre et al., 2017), the eyes (Lapan and Reddien, 2011; Lapan and Reddien, 2012; Mannini et al., 2004; Pineda et al., 2000), the central nervous system (Brown et al., 2018; Coronel-Córdoba et al., 2022; Cowles et al., 2013; Cowles et al., 2014; Currie and Pearson, 2013; Fraguas et al., 2014; März et al., 2013; Roberts-Galbraith et al., 2016; Ross et al., 2018), the epidermis (Dubey et al., 2022; Tu et al., 2015), the pharynx (Adler et al., 2014), the pigment cells (He et al., 2017; Wang et al., 2016), the musculature (Scimone et al., 2017; Scimone et al., 2018), the excretory system (Scimone et al., 2011), as well as for the establishment of axial polarity (Blassberg et al., 2013; Chen et al., 2013; Felix and Aboobaker, 2010; Hayashi et al., 2011; Pascual-Carreras et al., 2023; Scimone et al., 2014; Tian et al., 2021; Vásquez-Doorman and Petersen, 2014).

LIM-homeodomain (LIM-HD) proteins are a family of transcription factors that play a crucial role in cell fate specification, differentiation, and migration during embryonic development, especially for neural fates (reviewed in Hobert and Westphal, 2000; Kadrmas and Beckerle, 2004; Yasuoka and Taira, 2021). Structurally, LIM-HD proteins are characterized by the presence of two LIM domains, which mediates protein-protein interactions, and the homeodomain, which binds to specific DNA sequences and regulates gene expression (reviewed in Yasuoka and Taira, 2021). LIM domains and homeodomains are found in non-metazoan eukaryotes, but the specific combination of LIM-LIM-HD is only found in animals (Srivastava et al., 2010). During development of both vertebrate and invertebrate organisms, the expression of different combinations of *lim homeobox genes* (*lhx*) genes are considered to form a “LIM code” that specifies neural types within a tissue or organ and guide the establishment of topographically arranged connections (reviewed in Bachy et al., 2002; Gill, 2003; Hobert and Westphal, 2000; Shirasaki and Pfaff, 2002). For instance, during embryonic development of the mouse embryo, Lhx6 and Lhx7 specify GABAergic and cholinergic fates in cortical and forebrain neurons, respectively, (Lopes et al., 2012), while the axonal patterns, synaptic targets, and neurotransmitter profiles of dorsal hindbrain interneurons are instructed by Lhx1/5, Lmx1b and Lhx2/9 (Kohl et al., 2015). Moreover, LIM-HD have been involved in endodermal specification (Perea-Gómez et al., 1999; Satou et al., 2001; Wang et al., 2002), blastopore organizer activity (Yasuoka et al., 2009), head formation (Shawlot W et al., 1999) and in the regulation of the proliferation and migration of progenitor cells (Genead et al., 2010). Even though the important role of LIM-HD transcription factors during embryonic development has been well documented, our understanding about the role these genes may play during regeneration is more limited.

Previous studies have started to study the function of LIM-HD during planarian regeneration (Currie and Pearson, 2013; Forsthoefel et al., 2020; Hayashi et al., 2011; Li et al., 2019; März et al., 2013; Roberts-Galbraith et al., 2016; Scimone et al., 2014; Scimone et al., 2020). Here, we report the identification of the full LIM-HD repertoire in *Schmidtea mediterranea* that includes thirteen homologues belonging to the six evolutionary conserved types of *lhx* genes. We found that ten of them are associated with the central nervous system (CNS) and two with intestinal cells. A systematic functional RNAi analysis has uncovered Smed-LIM-HD functions in the specification of neural and intestinal cellular subtypes. We report that *Smed-islet1*, *-lhx1/5-1*, *-lhx2/9-3*, *-lhx6/8*, *-lmx1a/b-2* and *-lmx1a/b-3* are essential to pattern and size the planarian brain as well as for correct regeneration of specific subpopulations of dopaminergic, serotonergic, GABAergic and cholinergic neurons. *Smed-lhx1/5.2* and *Smed-lhx2/9.2* are required for the proper expression of diverse intestinal cell type markers, specifically the goblet subtype. Other LIM-HD are also involved in the control of axonal pathfinding (*lhx6/8*), axial patterning (*islet1* and *lmx1a/b-3*), head/body proportions (*islet2*) and stem cell proliferation (*lhx3/4, lhx2/9-3, lmx1a/b-2* and *lmx1a/b-3*) in planarians. Altogether, our results suggest that planarian LIM-HD could provide a combinatorial LIM code to specify distinct neuronal and intestinal cell identities during regeneration as well as to control axial patterning and axonal growing.

## 2. Results

### 2.1. Thirteen *lim homeobox* genes are present in *Schmidtea mediterranea*

Previous work had characterized 5 homologues of the LIM-Homeodomain (LIM-HD) family in planarians (Currie and Pearson, 2013; Forsthoefel et al., 2020; Hayashi et al., 2011; Li et al., 2019; März et al., 2013; Roberts-Galbraith et al., 2016; Scimone et al., 2014; Scimone et al., 2020). By *in silico* searches, we identified the presence of 13 *lim homeobox* genes (*lhx*) in the genome of *Schmidtea mediterranea*. All identified genes code for LIM-HD proteins that present the characteristic pair of LIM domains at the N-terminal and a Homeodomain at the C-terminal region. Phylogenetic analysis indicated that planarian possess representatives of all six major LIM-HD subfamilies (Figure S1). In particular, *S. mediterranea* possesses two *islet* genes, *Smed-islet1* (Hayashi et al., 2011; März et al., 2013) and *Smed-islet-2* (Li et al., 2019), two *lhx1/5* genes, named *Smed-lhx1/5-1* (Currie and Pearson, 2013) and *Smed-lhx1/5-1-2*, three members of the lhx2/9 gene subfamily, *Smed-lhx2/9-1* (Forsthoefel et al., 2020); *Smed-lhx2/9-2*; *Smed-lhx2/9-3*, a member for each of the *lhx3/4* and *lhx6/8* subfamilies, *Smed-lhx3/4* and *Smed-lhx6/8* (Roberts-Galbraith et al., 2016), respectively, and four homologues of the *lmx1a/b* subfamily, that were named *Smed-lmx1a/b-1*, *Smed-lmx1a/b-2*, *Smed-lmx1a/b-3*, *Smed-lmx1a/b-4*.

### 2.2. *lim homeobox* genes are mainly expressed in the planarian brain and digestive system

We sought to determine the expression pattern of all identified planarian *lim homeobox* genes by *in situ* hybridization in intact planarians (Figure 1) as well as by analyzing available Single-Cell (SC) transcriptomic data (Figure S2).

**Fig 1:**
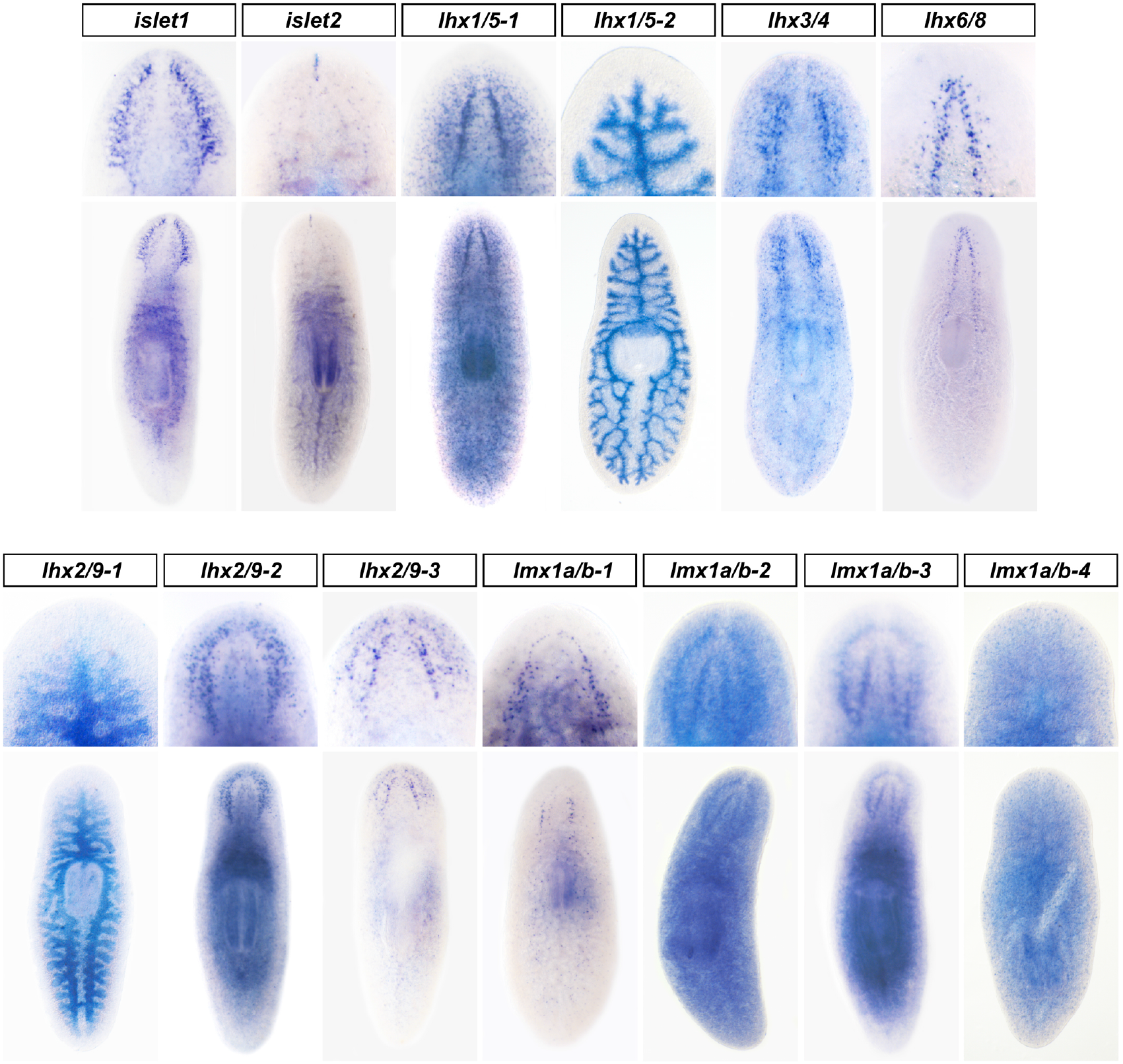
Planarian *lhx* are expressed in diverse domains of the cephalic ganglia and the digestive system. Representative expression patterns of planarian *lhx* genes in intact planarians as determined by whole mount *in situ* hybridization. The anterior end of the body is oriented towards the top. A magnification of the ventral view of the cephalic region or the dorsal view of the digestive system is shown on the upper panel for each of the genes analyzed.

Two planarian *lhx* genes, *lhx1/5-2* (this work) and *lhx2/9-1* (Forsthoefel et al., 2020) appeared strongly expressed in the planarian gut (Figure 1). The planarian intestine is a highly branched organ that consists in one anterior and two posterior primary gut branches that project into the head and tail, respectively, and that connects to a centrally located pharynx which evaginates ventrally through the mouth opening for feeding. Three distinct cell types have been reported within the planarian intestine: absorptive phagocytic cells, secretory goblet cells and basal cells, which locate in proximity to the basal region of the phagocytes (Fincher et al., 2018; Forsthoefel et al., 2020). According to available single cell data, *lhx1/5-2* is mainly expressed in differentiated phagocytes and in gut progenitor cells, while expression of *lhx2/9-1* is strongly enriched in differentiated basal and goblet cell types (Figure S2) (Fincher et al., 2018; Forsthoefel et al., 2020; Plass et al., 2018).

Notably, the expression of 10 out of the 13 planarian *lim homeobox* genes was enriched in the central nervous system (CNS) (Figure 1). The CNS of the planaria consists of a pair of cephalic ganglia that locate in the anterior region of the head and a pair of ventral nerve cords that run all along the anterior-posterior axis of the animal (Agata et al., 1998; Cebrià, 2007; Ross et al., 2017). Several studies have evidenced that the planarian brain is a complex organ that contain a large number of neural subtypes and that is regionalized in domains of gene expression along the anterior-posterior, dorsal-ventral and medio-lateral axis (Cebrià et al., 2002; Ross et al., 2017; Umesono et al., 1999). Interestingly, we identified that several *lhx* genes were expressed in distinct domains within the planarian brain (Figure 1). Previous studies reported *Smed-islet1* expression in the anterior and posterior poles during early stages of regeneration (Hayashi et al., 2011; März et al., 2013). We observed, in addition, strong expression of *Smed-islet1* in the planarian brain, particularly in the lateral brain branches, as well as in the parapharyngeal region of the body of intact animals. SC data confirmed the expression of *Smed-islet1* in differentiating secretory, cholinergic (*chat*+) and GABAergic neurons as well as in progenitor neural cells (Figure S2). We could also confirm the expression of *Smed-islet2* in the most anterior tip of the planarian head (Li et al., 2019). We also detected few scattered *islet2* expressing cells around the cephalic ganglia and throughout the planarian body as well as a broad domain of *islet2* expression in the pharynx and parapharyngeal region. These *islet2* expressing cells could correspond to differentiated secretory, cholinergic neurons (*chat*+) and epidermal cells according to SC data (Figure S2). In agreement with previous reports, our studies detected *Smed-lhx1/5-1* expression in a large number of discrete neural cells distributed throughout the body and that appeared particularly dense in the medial domain of the cephalic ganglia (Figure 1, Currie and Pearson, 2013). Previous reports have also shown *lhx1/5-1* expression in stem cells (Currie and Pearson, 2013). In addition, we observed strong *lhx1/5-1* expression in the pharynx, both by *in situ* and in SC transcriptomic data (Figure 1 and S2). Finally, the already reported expression of *Smed-lhx6/8 (arrowhead*) was also confirmed and detected in discrete cells that locate medially within the planarian brain (Roberts-Galbraith et al., 2016) and that could correspond to differentiated cholinergic and GABAergic neurons according to SC information (Figure S2).

The expression patterns of previously uncharacterized *lim homeobox* genes were also studied in detail. The expression of the single *lhx3/4* homologue was detected mainly in discrete neurons that appear predominantly concentrated in the most posterior region of the brain lobes, as well as in discrete cells located along the nerve cords and the planarian body edges (Figure 1). According to SC transcriptomic data these cells correspond to diverse differentiated cell types such as cholinergic and GABAergic neurons, muscular and parenchymal cells as well as to neural progenitor cells (Figure S2, Plass et al., 2018; Scimone et al., 2014). Two out of the three *lhx2/9* genes present in planarians also appeared expressed in the cephalic ganglia. Strong *lhx2/9-2* expression was detected in the brain, particularly in an external domain that could relate to the lateral brain branches, as well as in the parapharyngeal body region (Figure 1). According to SC transcriptomic data *lhx2/9-2* expressing cells relate to differentiated secretory, pharyngeal, muscular, and neural (*chat*+) cells, as well as to progenitors for the muscular and epidermal lineages (Figure S2). Similarly, the expression of *lhx2/9-3* was particularly enriched in the external domain of the cephalic ganglia that could relate to the lateral brain branches. We also found some scattered *lhx2/9-3* expressing cells distributed throughout the planarian body (Figure 1). According to SC information, these *lhx2/9-3* cells could correspond to differentiated muscular and neural (*chat*+, GABAergic) cells as well as to muscular and neural progenitor cells (Figure S2).

Finally, the expression of the four genes of the LMX1a/b subfamily was also characterized. Whole-mount *in situ* hybridizations revealed *lmx1a/b-1* expression in a discrete row of few neurons that seemed to trace the most anterior region of the brain commissure. Disperse *lmx1a/b-1* expressing cells were also found throughout the planarian body. Some of them appeared particularly condensed at the most posterior domain of the cephalic region. Those *lmx1a/b-1* expressing cells located ventral to the cephalic ganglia and might be defining a subdomain within the ventral nerve cords (Figure 1). In contrast to *lmx1a/b-1*, the expression of *lmx1a/b-2* and *lmx1a/b-3* appeared uniform and extended throughout the cephalic ganglia and the planarian body (Figure 1). *lmx1a/b-2* expression appeared relatively ubiquitous and SC transcriptomic data suggested that *lmx1a/b-2* transcripts are detected in differentiating neurons (*otf*+, *npp18*+) as well as in neoblasts and progenitor cells for the epidermal, muscular, intestinal and neural lineages, while *lmx1a/b-3* expression is also detected in several cell types, but mainly in differentiated protonephridia and neurons (*otf*+ *GABAergic*, *chat*+), as well as in epidermal, muscular, and neural progenitor cells (Figure S2). Finally, *Smed-lmx1a/b-4* expression was detected in discrete cells distributed all along the dorsal and ventral body. Available SC data suggest that *lmx1a/b-4* positive cells possess secretory (Figure S2, Plass et al., 2018) and/or neural identity (Fincher et al., 2018).

Our analyses reveal that *lhx* genes were broadly expressed in the planarian body, particularly in the neural, intestinal, pharyngeal, and muscular lineages, as well as in several stem cell progenitor subtypes. Moreover, the expression patterns observed for several *lhx* in subdomains along the anterior-posterior (*lhx3/4*) and medio-lateral (*islet1*, *lhx1/5-1*, *lhx6/8*, *lhx2/9-2* and *lhx2/9-3*) axes of the cephalic ganglia further evidence the molecular complexity of the planarian brain.

### 2.3. Planarian LIM-HDs control midline patterning and stem cell proliferation

To characterize the function of *Smed-lhx* genes we performed RNA interference (RNAi)-based functional analyses. We were interested in analyzing the regenerative process in the absence of *lhx* genes. Thereby, animals were amputated pre- and post-pharyngeally after dsRNA delivery and the regenerative process monitored for 10-14 days.

Apparent normal regeneration of the anterior blastemas was observed after silencing most of the planarian *lhx* genes (Figure 2A). However, the silencing of two *lhx* genes impaired anterior regeneration. Thus, RNAi against *islet1* and *lmx1a/b-3* resulted in midline defects and regeneration of merged eyes in n=53/62 and n= 9/50 of the analyzed animals, respectively. Our results for the silencing of *islet1* agree with previous reports (Hayashi et al., 2011; März et al., 2013). Also, in most conditions no obvious morphological defects were observed during posterior regeneration with the exception of the rounded and smaller blastemas observed in all *islet1* silenced animals (data not shown, see next sections).

**Fig 2.**
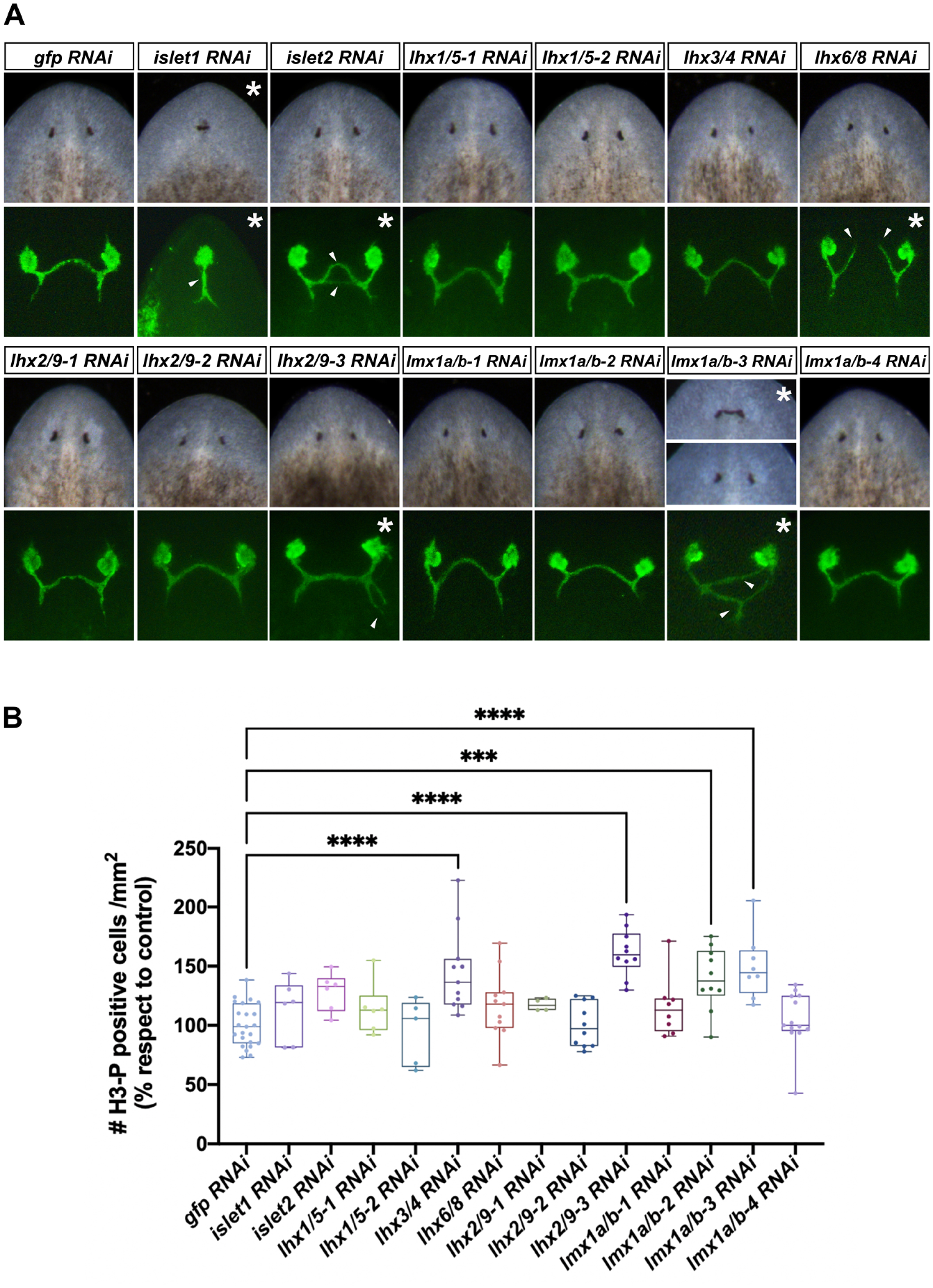
Defects in anterior regeneration and stem cell proliferation after *llhx* genes silencing. **A,** upper panels show live images of anterior facing regenerated blastemas. Bottom panels show regenerated eyes photoreceptor cells and their axonal projections as labeled with the VC1 antibody. All animals correspond to trunk pieces at 10-12 days of regeneration after 2 rounds of RNA interference. Asterisks highlight the panels where defects in the regenerative process are observed. White arrowheads in *islet1*; −2; *lhx6/8*; *lhx2/9-3* and *lmx1a/b-3* (RNAi) planarians point to aberrant eye photoreceptor axonal projections. **B**, Quantification of mitotic H3-P+ immunolabeled cells at 10 days of regeneration in control *gfp(RNAi*) and after silencing planarian *lhx* genes. Trunk pieces regenerating anterior and posterior wounds were analyzed and the total number of mitotic cells normalized to the body area. Between 4 and 20 animals were analyzed per RNAi condition. Values in graphs are represented as % respect to the mean of control *gfp* RNAi animals (*** p-value<0.001; **** p-value<0.0001).

To further characterize the regenerative process in the absence of *lhx* genes, we investigated the proliferative rate of stem cells and carried out immunostaining against a phosphorylated form of Histone-3 that identifies the G2/M stage of the cell cycle. We quantified the total number of mitoses in planarians regenerating anterior and posterior wounds at 10 days after amputation. In four of the RNAi conditions (*lhx3/4*, *lhx2/9-3*, *lmx1a/b-2* and *lmx1a/b-3*) we observed significantly increased rates of neoblast proliferation when compared to control *gfp*(RNAi) planarians (Figure 2B). Notably, all these four *lhx* genes were expressed in planarian stem cells (Figure S2, Scimone et al., 2014), suggesting that these genes could function autonomously in regulating planarian stem cell proliferative rates.

### 2.4. LIM-HDs are needed for proper visual axonal projections and brain regeneration

As we found that most of the *lhx* genes were expressed in the central nervous system (CNS) of the planarian, we investigated whether they were needed for the proper regeneration of the visual axons and the CNS. To do that we performed immunostainings against eye photoreceptor cells and with a panneural antibody.

Control *gfp*(RNAi) planarians regenerated eyes and visual axonal projections that connected stereotypically to the brain forming proper optic chiasms (Figure 2A, Sakai et al., 2000). We observed that abnormal visual axonal projections towards the planarian brain accompanied the midline defects and cyclopic eyes observed in silenced *islet1* and *lmx1a/b-3* animals (Figure 2A). Remarkably, axonal growth of the photoreceptor cells was also found perturbed in the apparently normal regenerated eyes of *islet2*, *lhx6/8*, *lhx2/9-3* and *lmx1a/b-3* silenced planarians, suggesting a role for those genes on axonal growth pathfinding (asterisks, Figure 2A). The most striking phenotype was observed after silencing *lhx6/8*. In those animals, the visual axonal projections completely failed to cross the midline and connect to the contralateral side. These results agree with previous data that reported a role of *lhx6/8* on defining neurons that serve as guidepost cells for *de novo* regeneration of the visual system (Roberts-Galbraith et al., 2016; Scimone et al., 2020).

Analyses of the nervous system of *lhx(*RNAi) animals visualized with the panneural marker anti-synapsin identified that the regeneration of the cephalic ganglia was altered after silencing several *lhx* genes (Figure 3). In particular, regeneration of fused or elongated brain ganglia occurred in *islet1* and *islet2* silenced planarians, respectively; moreover, RNAi of either *islet1, lhx1/5-1, lhx2/9-3* or *lmx1a/b-2* resulted in significantly smaller brains (Figure 3B). Finally, the silencing of *lhx6/8* disrupted the medial anterior commissure that connects both cephalic ganglia, in agreement of the observed visual axonal defects and previous reports (Roberts-Galbraith et al., 2016; Scimone et al., 2020).

**Fig 3:**
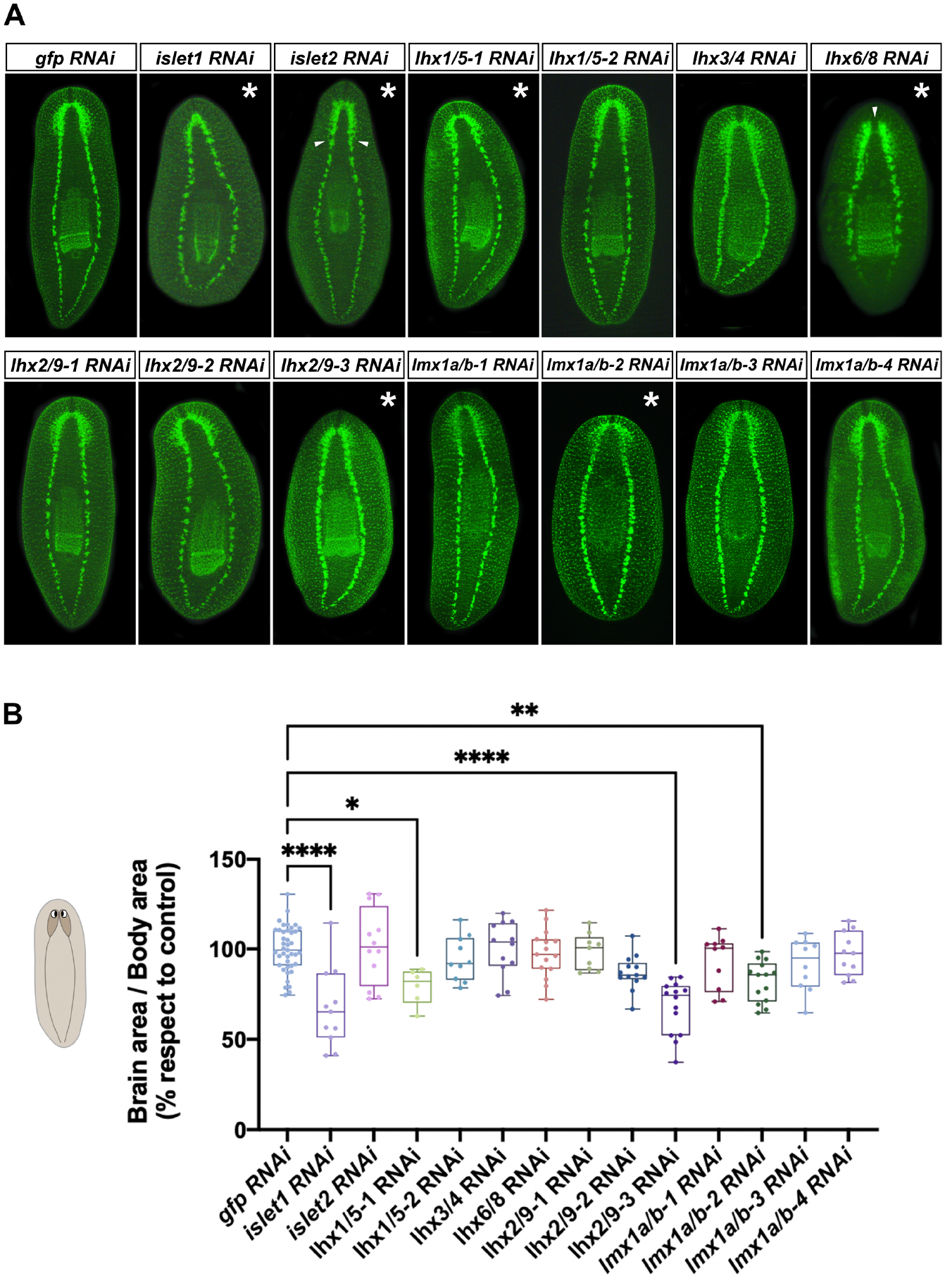
Planarian LIM-HD are required for proper brain regeneration. **A,** Planarian CNS as labeled with the anti-SYNAPSIN antibody. All animals correspond to trunk pieces at 10-12 days of regeneration after 2 rounds of RNA interference. Asterisks highlight the panels where defects in regeneration of the brain are detected. White arrowheads in *islet2(RNAi*) planarians point to brain ganglia that elongate posteriorly. White arrowhead in *lhx6/8* (RNAi) animals point to a medial gap in the brain commissure. **B,** Graphical representation of the ratio of the brain-to-body area in control *gfp(RNAi*) and after silencing planarian *lhx* genes. The brain and body area of planarians at 10-12 days of anterior and posterior regeneration were analyzed. Between 6 and 40 animals were analyzed per RNAi condition. Values in graphs are represented as % respect to the mean of control *gfp* RNAi animals (*p-value<0.05; **p-value<0.01; **** p-value<0.0001).

### 2.5. LIM-HDs specify distinct neural subtypes and are needed for correct patterning of the planarian brain

Regeneration of a functional brain requires the specification of several neural subtypes that need to be properly patterned and integrated into a functional unit. Therefore, we investigated next if planarian LIM-HD are involved in defining neural cell type identity during regeneration by analyzing the presence of five main neural subtypes after silencing each of the 13 *lhx* genes by RNAi (Figure 4 and Figure S4). We characterized the presence of dopaminergic neurons expressing *tyrosine hydroxylase* (*th*) (Nishimura et al., 2007a), octopaminergic neurons expressing *tyramine beta-hydroxylase (tbh*) (Nishimura et al., 2008b), GABAergic neurons expressing *glutamine decarboxylase (gad*) (Nishimura et al., 2008a), serotonergic neurons expressing tryptophan hydroxylase (*tph*) (Nishimura et al., 2007b) and cholinergic neurons expressing *choline acetyltransferase* (*chat*) (Nishimura et al., 2010).

**Fig 4:**
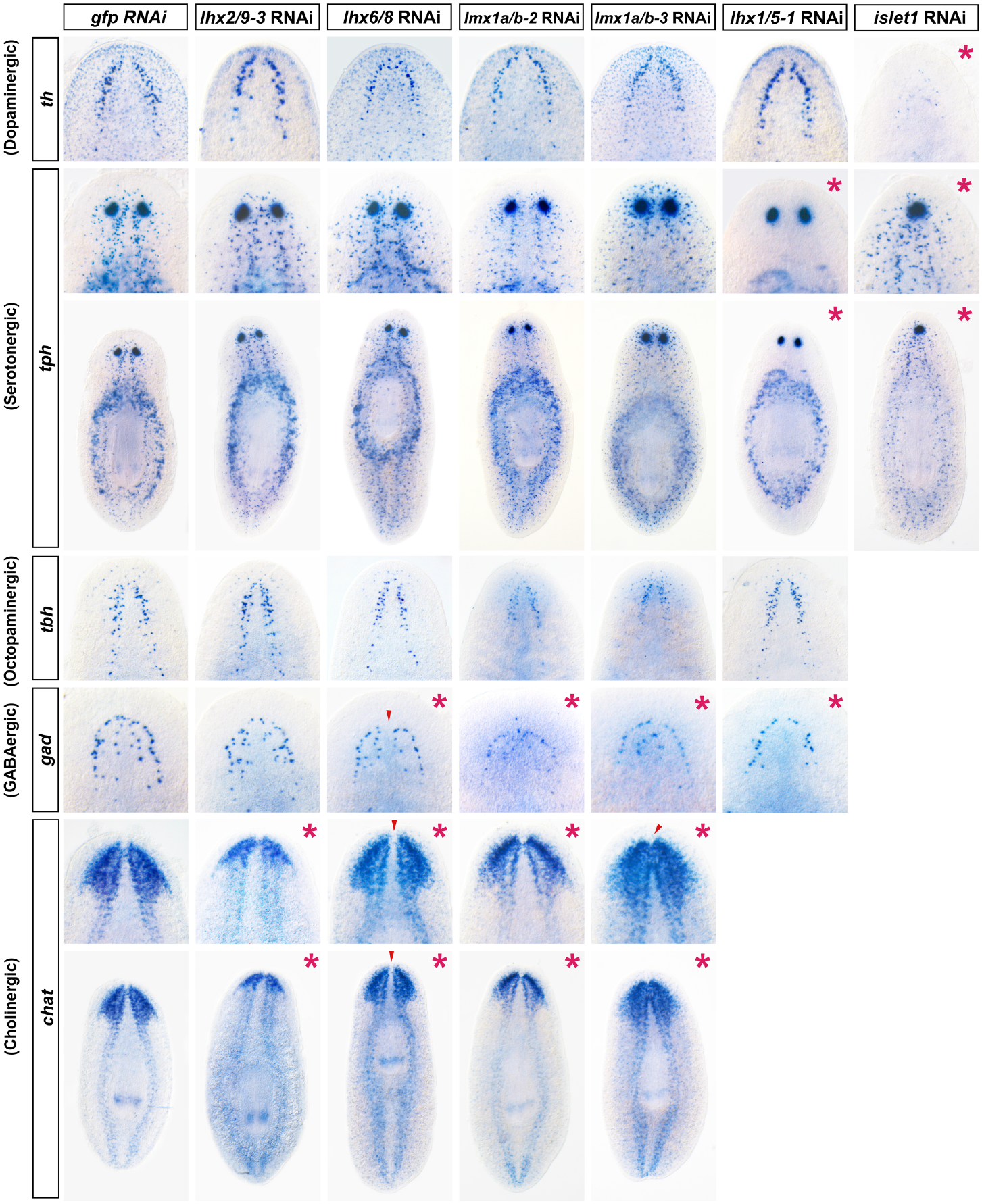
Aberrant regeneration of different neural cell subtypes after silencing planarian *lhx* genes. Whole mount *in situ* hybridizations for *th* (dopaminergic), *tbh* (octopaminergic), *gad* (GABAergic), *chat* (cholinergic) and *tph* (serotonergic) neural cell types in control *gfp*(RNAi) and after silencing of *lhx2/9-3, lhx6/8, lmx1a/b-2, lmx1a/b-3, lhx1/5-1* or *islet1* genes. All animals correspond to trunk pieces at 10-12 days of anterior-posterior regeneration after 2 rounds of RNA interference. Arrowheads in *lhx6/8* (RNAi) animals point to a medial gap in the brain commissure. Arrowheads in *lmx1a/b-3*(RNAi) point to a thicker brain commissure. Asterisks highlight the panels where defects are detected. Anterior to the top.

Silencing of either *islet1* or *lhx1/5-1* perturbed the expression of the serotonergic neuronal marker *tph*. Serotonergic *tph*+ neurons distributed along the planarian body were completely absent after silencing *lhx1/5-1* (as previously reported by Currie and Pearson, 2013), while *tph* expressing cells that locate around the pharynx and in the eye appeared unaffected. Notably, in contrast, we observed that *islet1*(RNAi) treatment decreased exclusively the parapharyngeal domain of expression of *tph* and did not perturb either scattered serotonergic *tph* expressing cells or eye *tph*+ expressing cells (Figure 4). Moreover, the silencing of *islet1* caused a strong reduction in the number of cells expressing the dopaminergic marker *th*; this neural subpopulation remained mainly unaffected after silencing any of the other *lhx* genes analyzed (Figure 4).

Regeneration of octopaminergic neurons as analyzed by the expression of *tbh* was not affected after RNAi of any of the *lhx* genes (Figure 4). Also, regeneration of GABAergic and cholinergic neurons occurred in all analyzed RNAis. However, several patterning defects as well as defective regeneration of some subdomains of expression for those neural subtypes were observed (Figure 4). We confirmed that *lhx1/5-1* (RNAi) worms lacked the ventral subpopulation of GABAergic cells (corresponding to the internal row of *gad*+ cells as observed from a dorsal view, (corresponding to the internal row of gad+ cells as observed from a dorsal view, Currie and Pearson, 2013), as well as the midline defects in *lhx6/8* (RNAi) treated animals which regenerated with a gap between the lobes of the cephalic ganglia (Figure 4, Roberts-Galbraith et al., 2016). In addition, we observed that GABAergic and cholinergic neuron regeneration was perturbed after silencing *lmx1a/b-2* and *lmx1a/b-3*. Silencing of *lmx1a/b-2* resulted in reduced *chat* expression in the internal domain of the brain lobes, suggesting the presence of a reduced number of cholinergic neurons on this region of the cephalic ganglia (Figure 4). Also, in agreement with the midline defects observed during regeneration of the planarian eyes, *chat* and *gad* expression in the anterior region of the cephalic ganglia appeared fused and thicker in *lmx1a/b-3* silenced worms. Finally, visualization of *chat* expressing cholinergic neurons confirmed that *lhx2/9-3* silenced planarians regenerated smaller brains compared to control *gfp*(RNAi) planarians (Figure 4).

Altogether, these data suggests that LIM-HD transcription factors are required for correct regeneration and patterning of the planarian brain as well as to define distinct neural identities, particularly for the serotonergic and the dopaminergic subtypes.

### 2.6. *lhx1/5-2* and *lhx2/9-2* silencing perturbs intestinal gene expression

We observed enriched expression of *lhx1/5-2* and *lhx2/9-1* in the planarian digestive system (Figure 1). As already mentioned, three main cell types constitute this highly branched organ: phagocytes, basal and goblet cells. Phagocytic cells can be identified by the expression of the *cathepsin La* marker (*ctsla*) (Forsthoefel et al., 2020). Basal cells can be visualized by the expression of the *solute carrier-family transporters 22 member 6 (slc22a6*) (Forsthoefel et al., 2020). Finally, secretory goblet cells located in the medial region of the intestine can be identified by the enriched expression of the metalloendopeptidase *cg7631* (Forsthoefel et al., 2020) as well as by the presence of the RAPUNZEL-1 protein (Reuter et al., 2015).

We analyzed the expression of *ctsla, slc22a6* and *cg7631* after *lhx1/5-2* and *lhx2/9-1* RNAi treatment to decipher whether these genes play a role in gut regeneration and intestinal cell type specification in planarians (Figure 5). Silencing of either *lhx1/5-2* or *lhx2/9-1* caused defects in the expression of intestinal markers but did not seem to severely affect branching and regeneration of the new gut. We observed that silencing of *lhx1/5-2* significantly reduced the expression of the markers analyzed for all three intestinal cell types, while *lhx2/9-1* silencing perturbed particularly the expression of markers for the basal and goblet cell subtypes (Figure 5A). Previous work reported a mild effect of *lhx2/9-1* silencing on goblet cell regeneration (Forsthoefel et al., 2020). Our data revealed a much stronger effect, as the expression of the goblet cell marker *cg7631* was particularly reduced in both *lhx1/5-2* and *lhx2/9-1* silenced planarians and corresponded to about 39% and 6% of the expression observed in control *gfp*(RNAi) treated animals (Figure 5B). Similarly, quantification of RPZ-1 expressing cells confirmed the strong reduction in the number of goblet cells that silencing *lhx1/5-2* or *lhx2/9-1* genes caused (Figure 5B).

**Fig 5:**
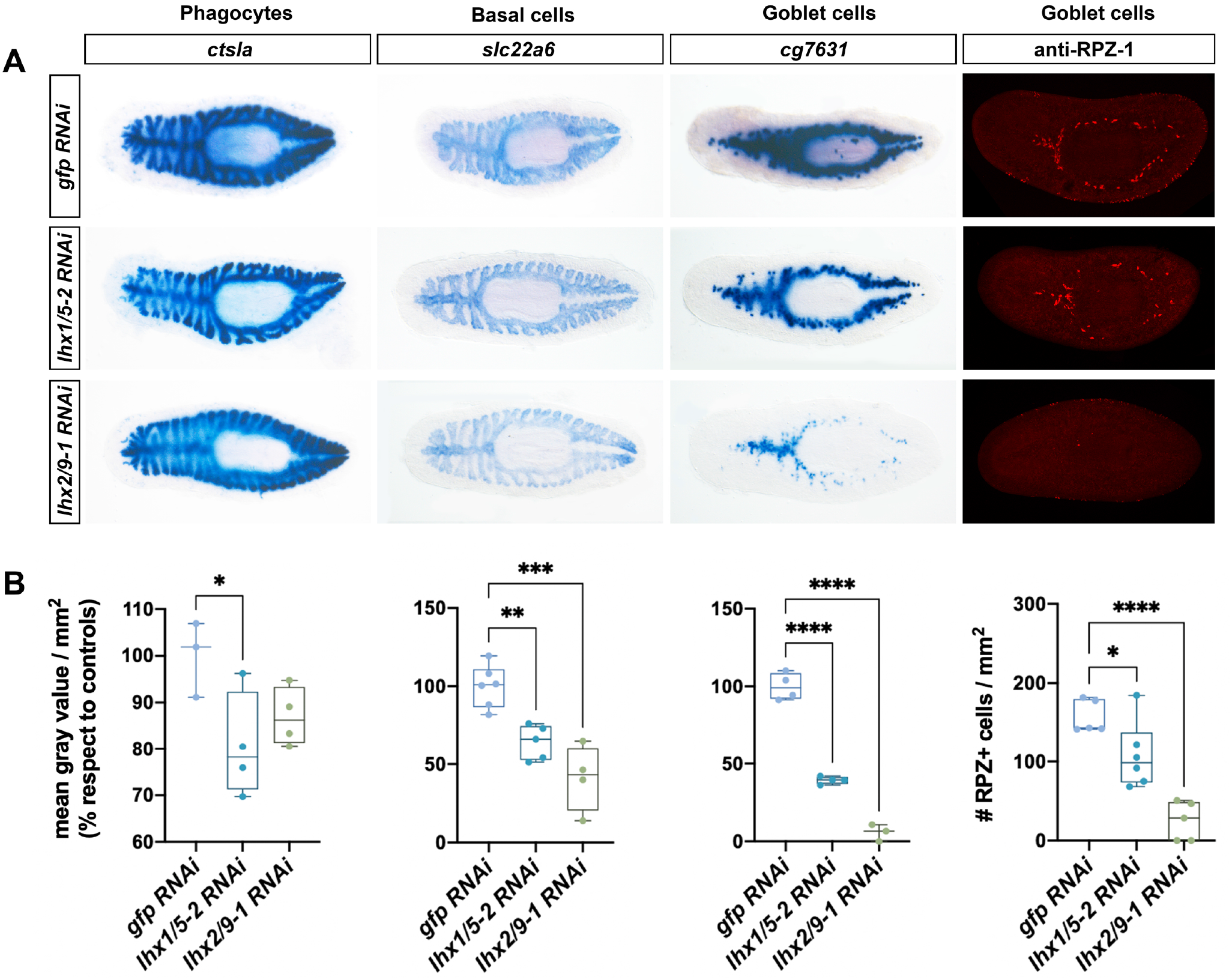
Defects in intestinal cell types after *Smed-lhx1/5-2* and *Smed-lhx2/9-2* silencing. **A,** Whole mount *in situ* hybridizations for *ctsla* (phagocytes), *slc22a6* (basal cells), *cg7631* (goblet) and immunostaining for RPZ-1 (goblet) intestinal cell types in control *gfp*(RNAi) and after silencing of *lhx1/5-2* or *lhx2/9-1* genes. All animals correspond to trunk pieces at 18 days of anterior-posterior regeneration after 2 rounds of RNA interference. Anterior to the left. **B,** Graphical representation of the quantification of the signal intensity of the colorimetric staining for the intestinal markers analyzed. Between 3 and 6 animals were analyzed per RNAi condition. Values in graphs are represented as % respect to the mean of control *gfp* RNAi animals, except for RPZ-1 where the values represent the quantification of the number of stained cells per body area. (*p-value<0.05; **p-value<0.01; *** p-value<0.0001; **** p-value<0.0001).

These data indicate that the expression of *lhx1/5-2* and *lhx2/9-1* is required for the proper expression of intestinal cell markers and the regeneration of the goblet cells.

### 2.7. Islet1 and Islet2 are needed for correct patterning and body proportions

During planarian regeneration, the expression of genes specific to anterior or posterior wounds is required for the specification of head and tail identities as well as for the differentiation of proper anterior and posterior structures (reviewed in Cebrià et al., 2018; Reddien, 2022). Expression of *islet1* in anterior and posterior blastemas during early stages of regeneration has been reported to be important for midline patterning and for the expression of *wnt1* and the establishment of posterior polarity (Hayashi et al., 2011; März et al., 2013). As expected, we observed that *islet1* (RNAi) treated planarians lacked posterior identity and the expression of *wnt1* in posterior wounds (Figure 6A), resulting in the regeneration of rounded posterior blastemas with fused nerve cords (Figure 6B). In addition, *notum* expression in anterior blastemas (Figure 6A) and regenerated eyes (Figure 2A) and cephalic ganglia (Figure 6B) merged at the midline after *islet1* silencing.

**Fig 6.**
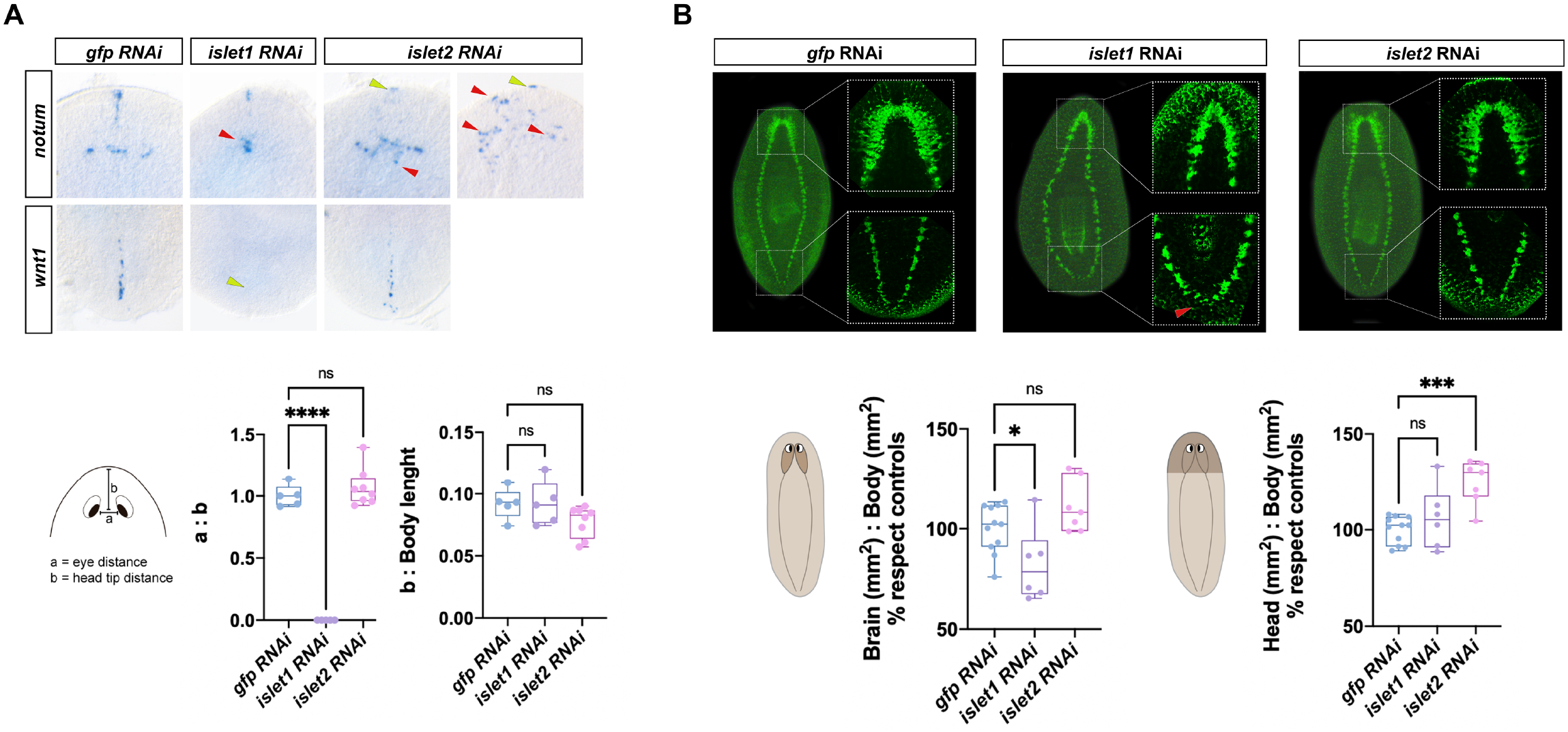
*islet1 (RNAi*) and *islet2* (RNAi) defects in patterning and body proportions. **A,** Whole mount *in situ* hybridizations for *notum* (in anterior blastemas) and *wnt1* (in posterior posterior) genes in control *gfp*(RNAi) and after silencing *islet* genes. Red arrowheads point to aberrant expression patterns. Yellow arrowheads point to absent expression. All animals correspond to trunk pieces at 10 days of anterior-posterior regeneration after 2 rounds of RNA interference. Anterior to the top. Graphical representation of both the eye distance to head tip distance and the head tip distance with respect to the body length. Between 5 and 8 animals were analyzed per RNAi condition. (**** p-value<0.0001). **B,** Immunostaining and confocal projections of the planarian CNS as labeled with the anti-synapsin antibody and graphical representation of the brain-to-body and the head-to-body ratios. Red arrowhead points to fused posterior nerve cords. Values in graphs are represented as % respect to the mean of control *gfp* RNAi animals. Between 6 and 11 animals were analyzed per RNAi condition. (*p-value<0.05; *** p-value<0.0001).

To clarify whether the expression of *islet2* in the anterior pole (Figure 1) could also be associated with the establishment of polarity, we analyzed the expression of genes associated with anterior (*notum*) (Petersen and Reddien, 2011) and posterior (*wnt1*) (Adell et al., 2009) identity in *islet2*(*RNAi*) treated animals. *notum* and *wnt1* were found expressed and restricted to anterior and posterior blastemas, respectively, suggesting that anterior and posterior identities were, probably, properly specified in *islet2* silenced planarians (Figure 6A). However, an unusual expression of *notum* was observed in the anterior blastema in *islet2*(RNAi) animals (Figure 6A). The normal expression of *notum* in the most anterior tip of the head region was barely detected in *islet2* silenced planarians; moreover, the *notum* expressing cells associated to the anterior brain commissure and the eye photoreceptors (Hill and Petersen, 2015; Scimone et al., 2020) appeared more scattered and abnormally distributed in this anterior region after *islet2(RNAi*) (Figure 6A).

The expression of *notum* in the head region has been associated with promoting brain size (Hill and Petersen, 2015) and facilitating and guiding the regeneration of the planarian visual system (Scimone et al., 2020). Therefore, we analyzed if the non-stereotypical pattern of *notum* expression observed in *islet2(RNAi*) planarians was linked to defects in brain size, eye positioning and/or visual axonal projection defects. As already mentioned, some aberrant projections of the visual axons toward the cephalic ganglia were observed (Figure 2A). Eye distance and eye positioning along the anterior-posterior axis of the planarian body were found to be normal in *islet2(RNAi*) worms (Figure 6A). Similarly, we found no differences in the size of the brain (Figure 6B) neither in the number of *cintillo*+ mechanosensory cells or *gad*+ GABAergic neurons (Figure S4), two population of neuronal cell types that have been strongly correlated with brain length and body size (Hill and Petersen, 2015; Oviedo et al., 2003; Takeda et al., 2009). On the other hand, quantification of the head-to-body ratio revealed that head regions were larger in *islet2*(RNAi) planarians compared to control *gfp*(RNAi) animals (Figure 6B). These results agree with the regeneration of normally sized but elongated brains after silencing *islet2* (Figure 3A).

Altogether, this data suggests that *islet2* is necessary for the stereotypical expression of *notum* in the head region, as well as for correct photoreceptor axonal projections and head-to-body allometric proportions.

## 3. Discussion

In this work we provide functional evidence that LIM-HD genes are involved in specifying neuronal identity in planarians, specially the dopaminergic and serotonergic neuronal subtypes, as well as in controlling the expression of intestinal markers and the regeneration of intestinal goblet cells. Our data confirms the previously reported role for *lhx1/5-1* in maintaining serotonergic and GABAergic neural cells (Currie and Pearson, 2013). In addition, we provide new evidence that *islet1* is necessary for the expression of *tph* in serotonergic parapharyngeal cells, as well as for the regeneration of *th*+ dopaminergic neurons. Also, our results suggest a role for *lmx1a/b-2* in cholinergic neurons. In both vertebrates and invertebrates, neural subtypes are specified by combinatorial expression of LIM-HD and other transcription factors; however, it is important to point out that the same neurons are not generated by the same combinations in different species (reviewed in Hobert and Westphal, 2000; Yasuoka and Taira, 2021). Previous studies in planarians have defined the role of several transcription factors in patterning the CNS and functionally specifying neural subtypes (reviewed in Ross et al., 2017). Thus, for instance, Nkx2.1 and Arx are required for the maintenance of cholinergic (ventral medial), GABAergic (ventral medial), and octopaminergic (correct number) neurons (Currie et al., 2016a); TCF1 is required for the regeneration of dorsal GABAergic neurons (Brown et al., 2018), and Pitx and Lhx1/5-1 play a role in maintaining the identity and function of serotonergic neurons in the planarian CNS (Currie and Pearson, 2013). Our results suggest that in planarians there might also be a combinatorial activity of LIM-HD and other transcription factors to correctly specify and/or pattern several neural subtypes such as serotonergic, dopaminergic, GABAergic and cholinergic during the regeneration of the CNS.

In the developing mammalian limb, Lim1 and Lmx1b function in controlling the initial trajectory of motor axons and establishing the fidelity of a binary choice (Kania et al., 2000). Similarly, in the fly, *lim3* participates in a combinatorial fashion with *tailup (Islet class*) to define the patterns of axonal projections of some motor neurons of the CNS (Thor et al., 1999). Previous works have reported the role of Lhx6/8 in promoting reconnection of the brain lobes and the visual system during planarian regeneration (Roberts-Galbraith et al., 2016; Scimone et al., 2020). Our data suggest that in addition to Lhx6/8, several other LIM-HD genes play a role in controlling the proper projections of the visual axons (Islet1, Islet2, Lhx2/9-3 and Lmx1a/b-3) and neural patterning (Islet1, Islet2, Lhx2/9-3, Lmx1a/b-2 and Lmx1a/b-3) during planarian regeneration.

In addition to the function of planarian LIM-HD in CNS patterning and neural specification, we report the role of two *lhx* genes (*lhx1/5-2* and *lhx2/9-1*) in the intestine. The current knowledge on the role of LIM-HD during intestinal development in other model systems is limited. It is known that Islet1 expression in the embryonic stomach activates Gata3 transcription to ensure normal pyloric development in the mouse (Li et al., 2014). Also, high levels of expression of Lhx1, Islet1, Islet2, and Lmx1a and the expression of Islet1 in stem-like cells have been reported in the mouse intestinal epithelium (Makarev and Gorivodsky, 2014). Lhx1 is also known for its role in blastoporal organizer activity during gastrulation (Yasuoka and Taira, 2021), as well as endodermal specification in ascidians (Satou et al., 2001), amphioxus (Wang et al., 2002) and mice (Perea-Gómez et al., 1999). In planarians, the activity of the transcription factor GATA4/5/6 is essential for the correct regeneration and maintenance of the gut (Flores et al., 2016; González-Sastre et al., 2017). A recent study has also uncovered intestine-enriched transcription factors that specifically regulate regeneration (hedgehog signaling effector gli-1) or maintenance (RREB2) of goblet cell types (Forsthoefel et al., 2020), as well as proposed a modest role for Lhx2/9-2 in maintenance of these planarian cell types (Forsthoefel et al., 2020). Here we further characterize Lhx2/9-2 as a major regulator of goblet cell regeneration. Even though we identified the expression of *lhx1/5-2* and *lhx2/9-1* in specific and different gut cell subtypes, we observed that the silencing of either of them strongly reduces the expression of markers for both goblet and basal cells. We also observed reduced expression of phagocyte markers when silencing *lhx1/5-2*. Further experiments might help to analyze whether these effects of the silencing of *lhx1/5-2* on different gut cell subtypes could be explained by its hypothetical expression in a presumptive common gut progenitor of the three main cell lineages. In any case, our results on the function of Lhx1/5-2 and Lhx2/9-1 on intestinal cells expands the limited current knowledge on the function of intestine-rich transcription factors in the regeneration of the planarian gut.

Amputation triggers planarian stem cell proliferation and promotes regeneration (Wenemoser and Reddien, 2010). Our results suggest that *lhx2/9-3, lhx3/4, lmx1a/b-2* and *lmx1a/b-3* limit the proliferation of the planarian neoblasts. Interestingly, all these genes are in part expressed by planarian stem cells and progenitor cell types (Plass et al., 2018; Scimone et al., 2014; Zeng et al., 2018), suggesting that they may play a cell autonomous function in controlling stem cell proliferation. These results agree with the reported role for *Lhx5* in the regulation of neural-precursor cell proliferation and migration during formation of the hippocampus in the mouse embryo (Zhao et al., 1999). In this model, precursor cells for the hippocampal anlagen are specified and proliferate in the absence of *Lhx5*, but many fail to exit the cell cycle (reviewed in Hobert and Westphal, 2000). Interestingly, the increased mitotic rates observed in *lhx2/9-3* or *lmx1a/b-2* silenced planarians are associated with regeneration of smaller brains, which could be caused by failure in exiting the cell cycle and in cell differentiation.

Finally, and in relation to the different abnormal phenotypes observed after silencing the several planarian *lhx* genes, it is worth mentioning the patterning defects observed after silencing *islet2*. As previously reported (Hayashi et al., 2011; März et al., 2013), we observed that planarian Islet1 is required for the correct establishment of the medio-lateral patterning and for the expression of *wnt1* in posterior wounds. Here, we report that a second islet gene, *Islet2*, is necessary for the stereotypical expression of *notum* in the head region, as well as to regenerate correct head-to-body allometric proportions. Considering the importance of *notum* for the establishment of anterior polarity as well as to regulate the size of the brain further studies should help to elucidate the relationship of *Islet2* and *notum*.

In other model systems, certain LIM-HDs function in combination in a well-defined ‘LIM code’. Thus, in the mouse developing forebrain, the differentiation of GABAergic neurons and cholinergic neurons is regulated by combinations of Lhx6, Lhx8, and Isl1(reviewed in Zhou et al., 2015); also, in the midbrain, cooperation of Lmx1a and Lmx1b regulate proliferation, specification, and differentiation of dopaminergic progenitors (Yan et al., 2011). Similarly, the Lhx1/5 genes *lin-11* and *mec-3* are both required for the terminal differentiation of a subset of specific motor neurons and interneurons in *C. elegans* (reviewed in Hobert and Westphal, 2000). In contrast to most invertebrate species that possess six *lhx* genes, one for each of the main LIM-HD subfamilies, we have identified several representatives of the *islet*, lhx1/5, *lhx2/9* and *lmx1a/b* classes. These genes have probably originated by internal duplications within the planarian lineage, as has been already reported for other gene families (Barberán et al., 2016b; Fraguas et al., 2021; Iglesias et al., 2008; Molina et al., 2009; Stelman et al., 2021; Su et al., 2017). Considering the established combinatorial role of LIM-HD genes in the control of neural identity during embryonic development, the coincident expression of some planarian *lhx* in some neural domains allow us to speculate that they might play a homologous role in specifying combinatorially the neural identify during regeneration of these animals. This is the case, for instance, of the domains of expression of *lmx1a/b-2* and *lmx1a/b-3, lhx1/5-1* and *lhx6/8*, or *lhx2/9-2* and *lhx2/9-3*. This hypothesis predicts that some *Smed-lhx* genes should be co-expressed in the same neuronal cells, which needs yet to be shown. Alternatively, the presence of a larger *lhx* repertoire in planarians could have also allowed to redefine specific functions for each of the LIM-HD in specifying distinct and unique cell types. Notably, unlike vertebrates, we have not observed a role for any of the planarian Lmx1a/b paralogs in specifying the dopaminergic neural subtype. We have identified, in contrast, that regeneration of these neurons depends on Islet1. Similarly, both *lhx2/9-2* and *lhx2/9-3* are expressed in lateral domains of the planarian brain, and *lhx1/5-1* and *lhx6/8* cells are detected in the medial region of the cephalic ganglia. Based on the coincidence of the territory of expression of these *lhx* genes in planarians, we cannot discard that the silencing of one of them is counteracted by the expression of the other paralog/paralogs. Combined RNAi would help to clarify, for instance, if like the mouse counterparts, planarian Lmx1a/b genes work together to regulate dopaminergic neurons or if only *islet1* play this role. These experiments would identify possible compensatory effects for those genes and further validate the presence of a LIM code in the worm.

In summary, here we report the full repertoire of *lhx* genes in planarians and describe their expression patterns at the levels of whole-mount and single-cell. RNAi functional analyses have uncovered novel functions for some of these genes mainly in the regeneration of specific neuronal and intestinal cell subtypes as well as on the proper patterning of the brain and body proportions.

## 4. Materials and methods

### 4.1. Animal culture

An asexual clonal line of the planarian species *Schmidtea mediterranea* was used for all experiments. Animals were maintained at 18-20°C in artificial planarian water as previously described (Cebrià and Newmark, 2005) and fed once a week with organic veal liver. All planarians were starved for at least 1 week before experiments.

### 4.2. Identification and isolation of *lim* homeobox genes

*lim homeobox* genes were identified from the genome of *S. mediterranea* (Grohme et al., 2018) using the blast tool of Planmine v3.0 (Rozanski et al., 2019). The protein sequence of LIM-HD protein homologs of humans, flies, and planarians were used as queries. The protein domain conservation of the planarian candidate transcripts was analyzed using the SMART (http://smart.embl-heidelberg.de) and Pfam protein domain databases (http://pfam.xfam.org/). The presence of two LIM domains in the amino termini and a centrally located HD was used as a condition to select the candidates. TRIzol® reagent was used to extract total RNA from a mix of regenerating and intact planarians, and cDNA was synthesized with Superscript III® following manufacturer’s instructions. All identified *lim homeobox* genes were amplified using specific primers (Table S1). PCR products were cloned into PCRII® vectors for synthesis of ssRNA-DIG labeled probes. Synthesis of dsRNA for RNA interference experiments was performed by incorporating T7 and SP6 sequences to the PCR products.

### 4.3. Phylogenetic analyses

Protein sequences of LIM-HD homologues were obtained from NCBI and Planmine v3.0 (Rozanski et al., 2019) and aligned using MAFFT with the L-INS-i strategy (Katoh et al., 2018). The aligned full sequence was used to reconstruct the phylogenetic tree. The phylogenetic tree was inferred with the IQ-TREE web server, with default options, including the automatic substitution model selector, the ultrafast bootstrap analysis (1000 replicates) and the single branch test number (1000 replicates) (Minh et al., 2013; Trifinopoulos et al., 2016). The approximate Bayes test option was selected. The phylogenetic tree was visualized using iTOL (https://itol.embl.de) and edited with Adobe Illustrator. Accession numbers for planarian *Schmidtea mediterranea* transcripts retrieved from Planmine can be found in Supplementary Table 1. Accession numbers of LIM-HD protein sequences retrieved from NCBI are as follows: *Drosophila melanogaster* (*Dme*) Lim1A NP_572505.1, Lim3 NP_001260559.1, Lmx1a NP_729801.1, Dme_CG4328 NP_648567.2, Apterous NP_001163058.1, Arrowhead NP_001261379.1, Tailup NP_476774.1; *Homo sapiens* (Hsa) Islet1 NP_002193.2, Islet2 EAW99220.1, Lhx1 NP_005559.2, Lhx2 NP_004780.3, Lhx3 AAG10399.1, Lhx4 NP_203129.1, Lhx5 NP_071758.1, Lhx6 NP_055183.2, Lhx8 AAH40321.1, Lhx9 NP_064589.2, Lmx1b NP_796372.1, Lmx1b AAI43802.1; *Mus musculus* (*Mmu*) Islet1 EDL18368.1, Islet2 EDL25854.1, Lhx1 NP_032524.1, Lhx2 NP_034840.1, Lhx3 AAI50690.1, Lhx4 NP_034842.2, Lhx5 NP_032525.1, Lhx6 CAA04011.1, Lhx8 NP_034843.2, Lhx9 NP_001036042.1, Lmx1a XP_030097992.1, Lmx1b XP_006497809.1; *Nematostella vectensis* (Nve) Islet1 XP_032218609.1, lhx1 XP_001631723.2, lhx3/4 XP_048587815.1, Awh XP_032229933.1, lhx2/9 XP_032240842.1, lmx1 XP_048587096.1; *Strongylocentrotus purpuratus* (Spu) Islet1 XP_781774.3, Lim1 NP_999810.1, Lim1b XP_030853804.1, Lhx2/9 XP_030853437.1, Lhx2/9-like XP_030853097.1, Lhx3/4 XP_030852869.1, Lhx6/8_Awh XP_030853744.1.

### 4.4. Whole-mount in situ hybridization

Colorimetric whole-mount *in situ* hybridization (WISH) was performed as previously described (Currie et al., 2016b). Animals were sacrificed by immersion in 5% N-acetyl-L-cysteine (5 minutes), fixed with 4% formaldehyde (15 minutes) and permeabilized with Reduction Solution (10 minutes). Riboprobes were synthesized using the DIG-labelling kit (Sp6/T7) from Roche. Animals were mounted in 70% glycerol before imaging.

### 4.5. Single-cell expression profiles

The single-cell sequencing data expression profiles of *lim homeobox* genes during cell differentiation (Plass et al., 2018) and in neoblasts (Zeng et al., 2018)were obtained from Planmine (Rozanski et al., 2019).

### 4.6. RNA interference (RNAi)

Double-stranded RNA (dsRNA) was synthesized by *in vitro* transcription (Sp6/T7 from Roche) as previously described (Sánchez-Alvarado and Newmark, 1999). Two rounds of RNAi were performed for all *lim homeobox* genes. In each round, dsRNA was injected into the digestive system of each planarian during three consecutive days. Three doses of 32.2 nl (at 1 mg/mL) were delivered each day using a Nanoject II (Drummond Scientific Broomall, PA, USA). The head and tail of the animal was amputated on the fourth day. Planarians trunk pieces were allowed to regenerate for 4 days before starting the second round of dsRNA injection and amputation. Control animals were injected with *gfp* dsRNA. Regeneration of the treated animals was observed for 10-14 days before proceeding for WISH and immunohistochemistry experiments.

### 4.7. Immunohistochemistry staining

Whole-mount immunohistochemistry was performed as previously described (Fraguas et al., 2021; Ross et al., 2015). The following antibodies were used: mouse anti-SYNAPSIN, used as pan-neural marker (anti-SYNORF1, Developmental Studies Hybridoma Bank, Iowa City, IA, USA) diluted 1:50; mouse anti-VC1 (Sakai et al., 2000), specific for planarian photosensitive cells (anti-arrestin, kindly provided by H. Orii and Professor K. Watanabe) diluted 1:15000; rabbit anti-phospho-histone H3 (Ser10) to detect cells at the G2/M phase of cell cycle (H3P, Cell Signaling Technology) diluted 1:300; anti-RAPUNZEL-1 (Reuter et al., 2015), used as a marker for intestinal goblet cells (RPZ-1, kindly provided by K. Bartscherer) used at 1:200. The secondary antibodies Alexa 488-conjugated goat anti-mouse and Alexa-568-conjugated goat anti-rabbit (Molecular Probes, Waltham, MA, USA) were diluted 1:400 and 1:1000, respectively. Samples were mounted in 70% glycerol before imaging.

### 4.8. Microscopy, image processing and quantification

Live animals were photographed with an sCM EX-3 high end digital microscope camera (DC.3000s, Visual Inspection Technology). Fixed and stained animals were observed with a Leica MZ16F stereomicroscope and imaged with a ProgRes C3 camera (Jenoptik, Jena, TH, Germany). Confocal images were obtained with a Zeiss LSM 880 confocal microscope (Zeiss, Oberkochen, Germany). Image processing and quantifications were performed with Adobe Photoshop and ImageJ2. Counting of the H3P-positive (Figure 2B) and RPZ-1 positive cells (Figure 5) was carried out manually and normalized by the total body area. Brain (Figure 3B) and head (Figure 6) areas were measured in anti-SYNORF1 and DAPI stained planarians and normalized by the total body area. Signal intensity quantification of colorimetric WISH (Figure 5) was done as previously described (Rossi et al., 2018): in all experimental conditions compared, animals were developed, stopped, and processed in parallel and for the same time; each single animal was photographed at the same magnification and exposition, and similarly processed with ImageJ2; images were converted in grayscale mode and a threshold was set manually (the same for all images of a given marker); the mean gray value was measured for each image and normalized to the area of the animal; values in graphs are represented as % respect to the mean of control *gfp* RNAi animals.

### 4.9. Statistical analyses and graphical representation

Statistical analyses and graphical representations were performed using GraphPad Prism 9. One-way ANOVA was performed to compare the means between conditions after confirming data normality and homogeneity using the Shapiro-Wilk test.

## Acknowledgments

We thank all members of the F. Cebrià, E. Saló and T. Adell laboratories for discussions, H. Orii and Prof. K. Watanabe for providing anti-VC-1 and K. Bartscherer for providing the anti-RPZ-1 antibody. Monoclonal anti-SYNORF1 antibody was obtained from the Developmental Studies Hybridoma Bank, developed under the auspices of the National Institute of Child Health and Human Development and maintained by the Department of Biological Sciences, University of Iowa, Iowa City, IA, USA.

## Author contributions

Conceived and designed the experiments, M.D.M, F.C; performed the experiments, M.D.M, D.A., S.F.; analyzed the data, M.D.M; wrote the paper, M.D.M, F.C, Funding acquisition, M.D.M, F.C.

## Figure legends

**FigS1. Phylogenetic tree of LIM-HD proteins.** Evolutionary relationship of Smed-LIM-HDs and other metazoan LIM-HD proteins reconstructed using the maximum likelihood method. The aligned full sequence of LIM-HD proteins was used to reconstruct the phylogenetic tree. Smed-LIM-HDs are highlighted with a colored star. The amino acid positions used for each protein are indicated after the species name. Support values written on the legend correspond to SH-aLRT (%) / aBayes / ultrafast bootstrap (%) support. Only the values of certain branches are indicated. The scale indicates expected amino acid substitutions per site. The tree was rooted with LMO (LIM Only) protein homologues.

**Fig S2. Graphical representation of *lhx* gene expression.** Schematic representation of the single-cell transcriptomic expression of *Smed-lim homeobox* genes in planarian differentiating cells (according to Plass et al., 2018) and neoblast progenitors for the diverse cellular lineages (according to Zeng et al., 2018). Single-cell data reveals the expression of *Smed-islet1* in differentiating secretory, cholinergic (*chat*+) and GABAergic neurons as well as in progenitor neural cells. *Smed-islet2* transcripts are mainly present in differentiated secretory, cholinergic neurons (*chat*+) and epidermal cells. *lhx1/5-1* expression is enriched in differentiated otf 2+ cells. *lhx1/5-2* is mainly expressed in differentiated phagocytes and in gut progenitor cells. The expression of *lhx3/4* is detected in differentiated neurons (*chat*+ and *npp18+*) and muscular cells and in neural progenitor cells. *lhx6/8* is detected in differentiated neurons (*chat*+ *cav-1*+, GABAergic) and secretory cells. *lhx2/9-1* expression is enriched in goblet cells. *lhx2/9-2* expression is mainly detected in differentiated secretory, pharyngeal, muscular and neural (*chat*+) cells, as well as in some progenitors for the muscular and epidermal lineages. *lhx2/9-3* transcripts are detected in differentiated muscular, neural (*chat*+, GABAergic) cells and in muscular and neural progenitors. *lmx1a/b-1* is mainly expressed in differentiated neurons (*chat*+, *cav-1*+, *otf*+). *lmx1a/b-2* is expressed quite ubiquitously and their transcripts seem particularly enriched in differentiating neurons (*otf*+, *npp18*+) as well as in neoblasts and progenitor cells for the epidermal. muscular, intestinal and neural lineages. *lmx1a/b-3* is detected in several and diverse cell types, mainly in differentiated protonephridia and neurons (*npp18*+, *otf*+, *GABAergic*, *chat*+), as well as in epidermal, muscular, and neural progenitor cells. *lmx1a/b-4* is enriched in differentiated secretory cells.

**Fig S3. Correct neural cell subtypes regeneration after silencing *planarian lhx* genes.** Whole mount *in situ* hybridizations for *th* (dopaminergic), *tbh* (octopaminergic), *gad* (GABAergic), *chat* (cholinergic) and *tph* (serotonergic) neural cell types in control *gfp*(RNAi) and after silencing of *islet2, lhx1/5-2, lhx2/9-1, lhx2/9-2, lhx3/4, lmx1a/b-1 or lmx1a/b-4* genes. Regeneration of all neural subtypes analyzed was not affected in either RNAi condition. All animals correspond to trunk pieces at 10-12 days of anterior-posterior regeneration after 2 rounds of RNA interference. Anterior to the top.

**Fig S4. Planarians regenerate a correct number of *cintillo* and *gad* expressing cells in the absence of *islet2*.** Expression of *cintillo* (chemosensory neurons), *gad* (GABAergic neurons) and *gpas* (lateral brain branches) in control *gfp* (RNAi) and *islet2* (RNAi) planarians by *in situ* hybridization. Quantification of the number of *cintillo* and *gad* expressing cells per body length reveals no significant (ns) differences between *gfp* (RNAi) and *islet2* (RNAi) treated planarians. All animals correspond to trunk pieces at 10-12 days of anterior-posterior regeneration after 2 rounds of RNA interference. Anterior to the top.

**Table S1. Planarian *lhx* sequence information.** Planarian *lim homeobox* genes transcripts ID and sequence, primers used for synthesis of ssRNA-riboprobes and for synthesis of dsRNA for RNA interference experiments.

