## Supplementary figures and images for "LIM-HD transcription factors are required for regeneration of neuronal and intestinal cell subtypes in planarians"

### Supplementary Figures

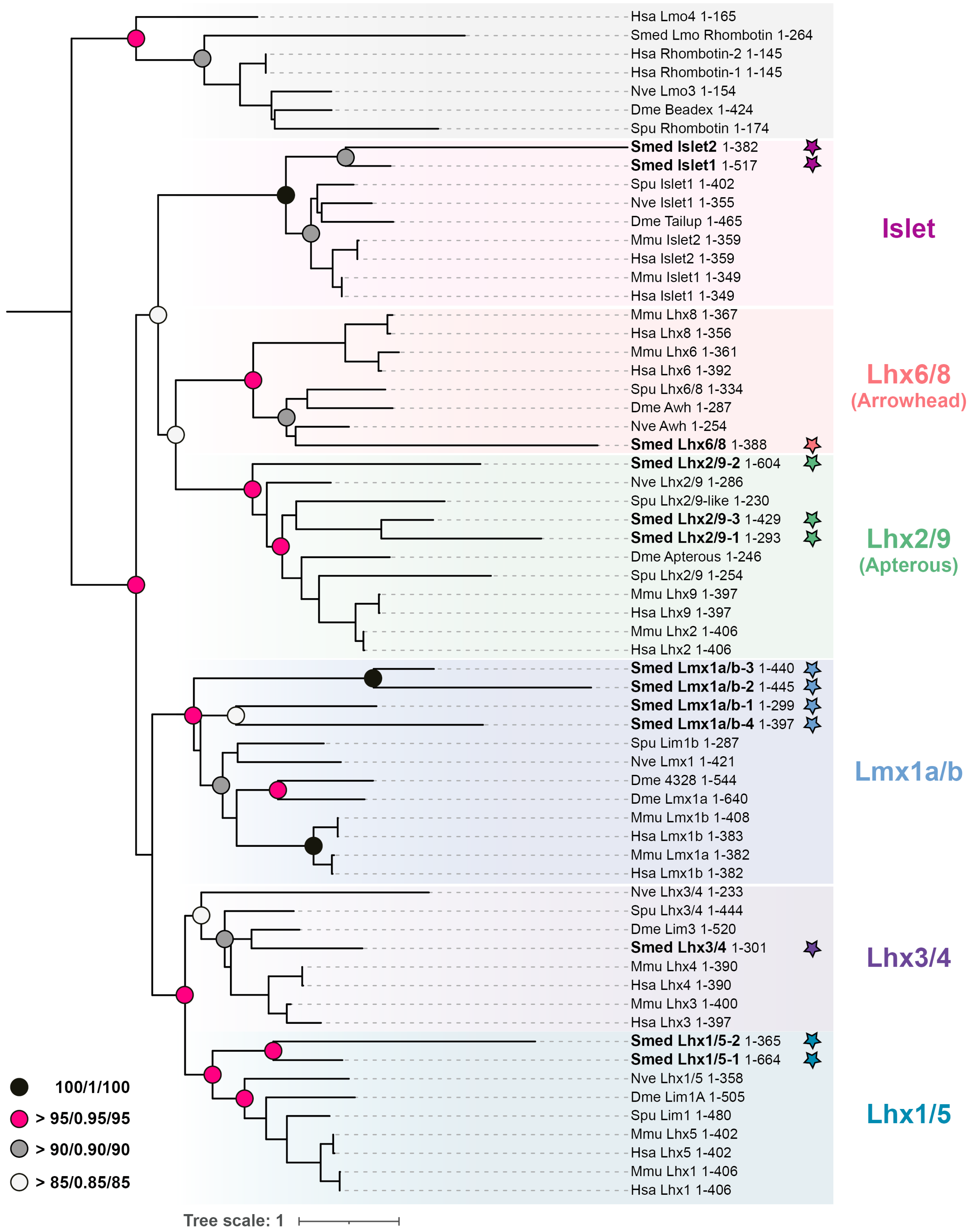

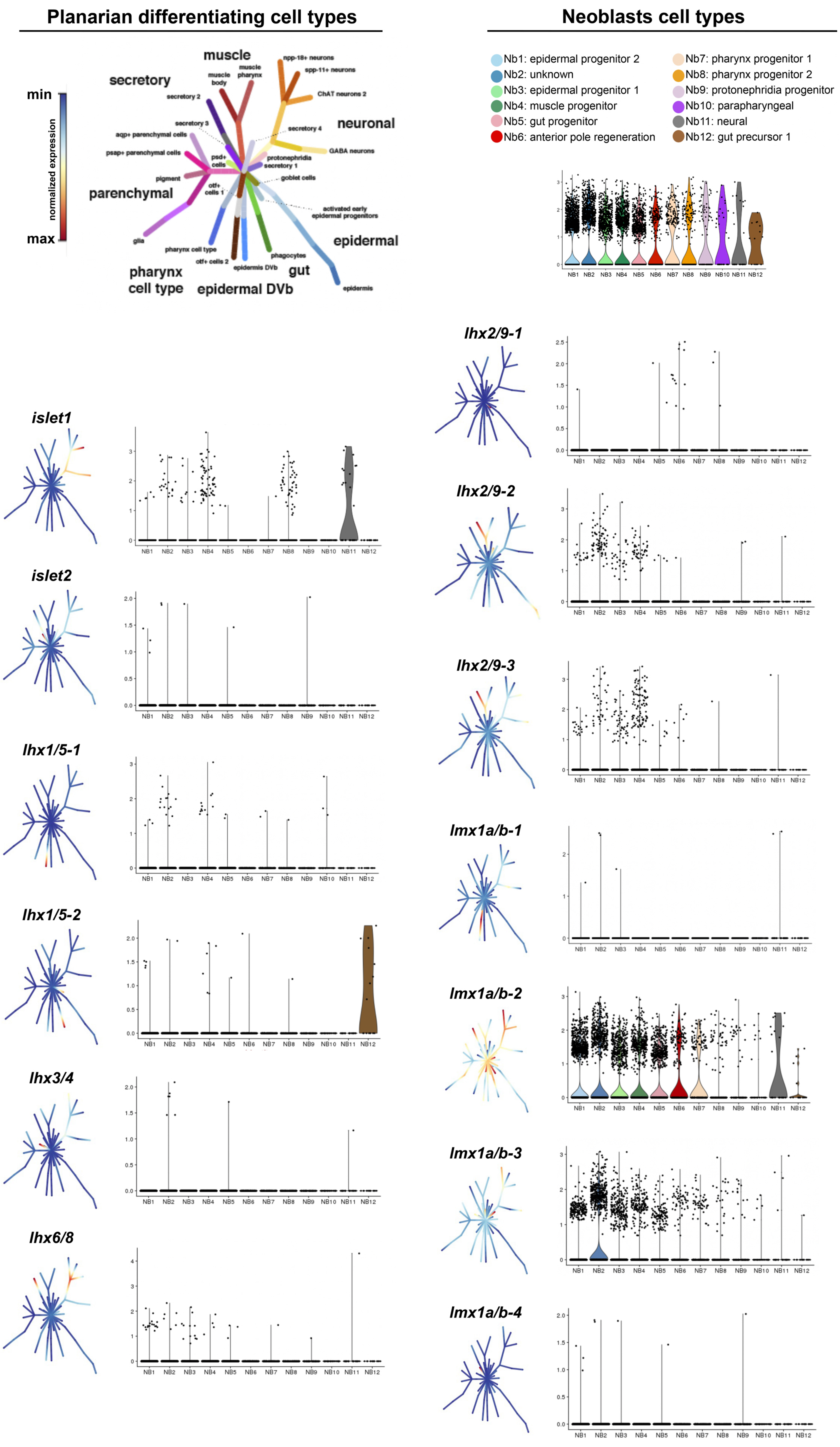

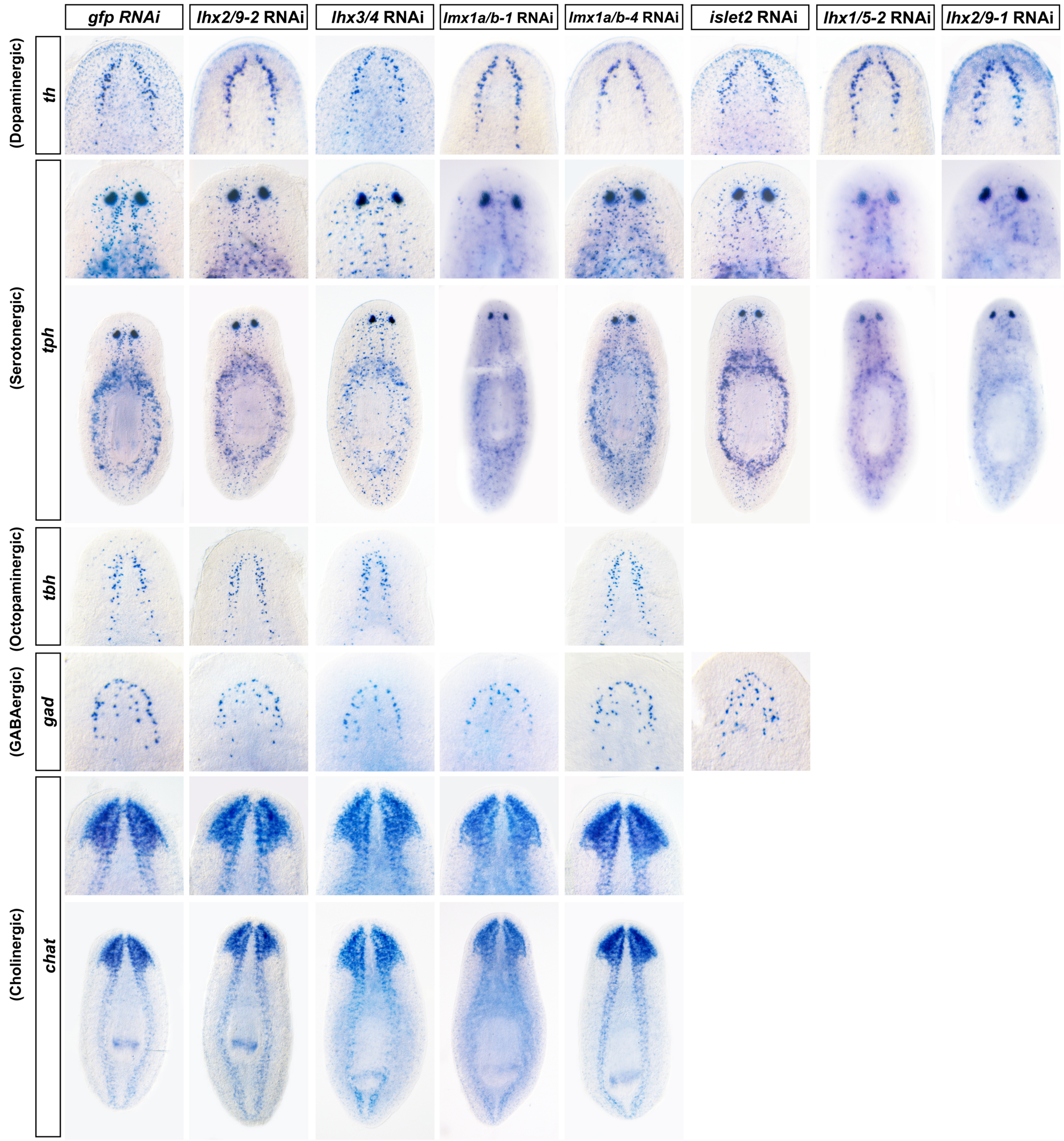

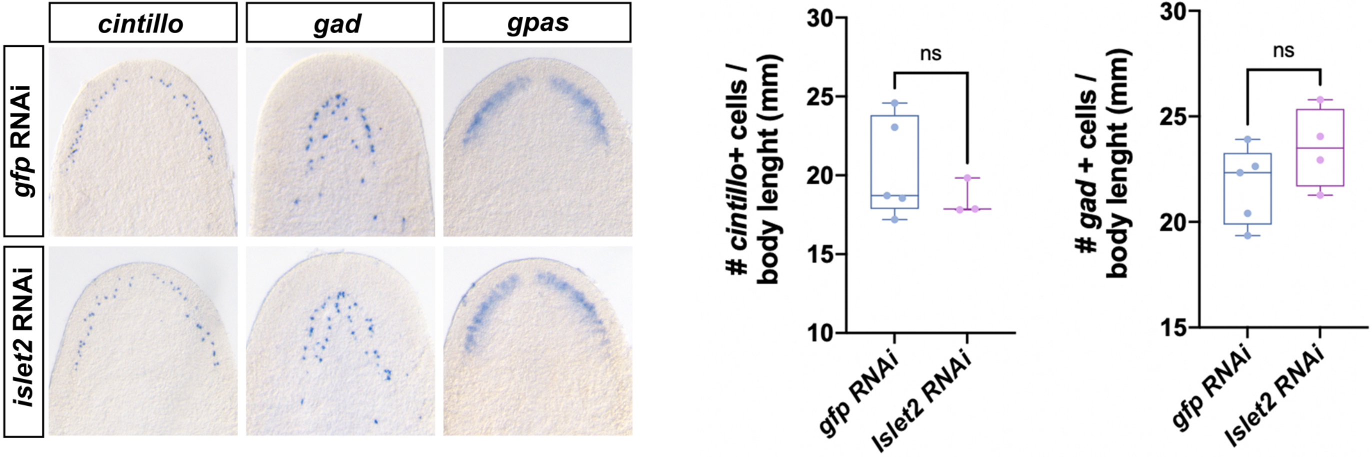
